# Presence of a D2 receptor truncated protein in the brain of the D2 receptor functional knockout mouse, revisiting its molecular and neurochemical features

**DOI:** 10.1101/757609

**Authors:** Natalia Sánchez, Montserrat Olivares-Costa, Marcela P González, Angélica P Escobar, Rodrigo Meza, Roberto Munita, Mauricio Herrera-Rojas, María E Andrés

**Affiliations:** Department of Cellular and Molecular Biology, Faculty of Biological Sciences, Pontificia Universidad Católica de Chile. Santiago 8331150, Chile; Department of Anatomy, Faculty of Medicine, Pontificia Universidad Católica de Chile. Santiago 8331150, Chile

## Abstract

Null mice for the dopamine D2 receptor (D2R) have been instrumental in understanding the function of this protein in the central nervous system. Several lines of D2R knockout mice have been generated, which share some characteristics but differ in others. The D2R functional knockout mouse, first described in 1997, is functionally null for D2R-mediated signaling but the Drd2 gene was interrupted at the most extreme distal end leaving open the question about whether transcript and protein are produced. We decided to determine if there are D2R transcripts, the characteristics of these transcripts and whether they are translated in the brain of D2R functional knockout mice. Sequence analysis of 3’ Rapid Amplification of cDNA Ends showed that D2R functional knockout mice express transcripts that lack only the exon eight. Immunofluorescence showed D2R-like protein in the brain of the knockout mice. As previously reported, D2R functional knockout mice are hypoactive and insensitive to the D2R agonist quinpirole (QNP). However, the heterozygous showed locomotor activity and response to QNP similar to the wild-type mice. Intriguingly, microdialysis experiments showed that heterozygous mice, such as knockouts, have half the normal levels of synaptic dopamine in the striatum. However, heterozygous mice responded similarly to wild-type mice to an acute injection of QNP, showing a 50% decrease in synaptic dopamine. In conclusion, D2R functional knockout mice express transcripts that lead to a truncated D2R protein that lacks from the sixth transmembrane domain to the C-terminal end but retains the third intracellular loop. We discuss the implications of this truncated D2R coexisting with the native D2R that may explain the unexpected outcomes observed in the heterozygous. Finally, we suggest that the D2R functional knockout mouse can be a useful model for studying protein-protein interaction and trafficking of D2R.

## Introduction

Dopamine is a neurotransmitter that participates in the control of voluntary movements and motivated behaviors, among other relevant functions. Two types of receptors mediate the action of dopamine, type 1 receptors coupled to excitatory G protein and which include D1 and D5, and type 2 receptors coupled to inhibitory G protein, which include D2, D3 and D4 receptors. D2 receptors (D2R) exist as two splice variants, the long (D2L) and the short (D2S) variants that differ in 29 amino acids in the third intracellular loop [1, 2]. Cumulated evidence indicate that D2L variant mediates mainly dopamine postsynaptic actions whereas D2S variant mediates presynaptic control of dopamine release and dopamine neurons firing [3]. Although, more recently it was described that under basal conditions both isoforms are able to play postsynaptic functions [4].

D2R are especially abundant in the striato-pallidal efferent GABA medium spiny neurons (MSN) that are involved in the control of voluntary movements [5, 6]. The activation of postsynaptic D2R in these GABA efferent MSN induces locomotor activity by decreasing the inhibitory action that this pathway, called “indirect pathway” have on locomotion [7]. Besides, D2R are present in presynaptic dopamine and glutamate afferents to the striatum [8], where negatively regulate the release of these neurotransmitters [8, 9]. D2R localized in the soma of dopamine neurons also regulate the firing of these neurons. Activation of D2R either in the soma or terminals of dopamine neurons will decrease synaptic dopamine in the striatum, which in turns will decrease locomotor activity by increasing the action of D2R-driven indirect pathway.

Different D2R deficient mice have been generated, which have served to reveal the multiple roles of this receptor [10, 11, 12]. The D2R knockout (KO) mice, generated by Baik et al. (1995) and by Jung et al. (1999), have a deletion of the exon 2, producing a mouse with a total deficiency of D2R protein. On another hand, Kelly et al. (1997) interrupted the *Drd2* gene in the most distal part of the gene and proved that these mice lack binding of D2R ligands.

Interested in the D2R functional knockout mouse, when characterizing it, we observed that this mouse expresses D2R mRNA. Thus, to determine why the mRNA is produced, we investigated the type of deletion made in the Drd2 gene using current genome information. Also, we sequenced the transcripts and searched for the protein in the brain of D2R functional knockout mice. Moreover, we compared behavioral and neurochemical features in the three genotypes. We found that the D2R functional KO mouse expresses several transcripts that have in common the lack only of the exon 8, leading to a truncated D2R protein that loses the sixth and seventh transmembrane domain and the C-terminal end. Immunofluorescent studies showed the presence of D2R-like proteins in the brain of the D2R functional knockout mice. Therefore, we propose that the D2R functional knockout mouse is a good model to study D2R protein interaction and trafficking.

## Materials and Methods

### Animals

Male and female C57BL/6 heterozygous D2R mice [11] were kindly donated by Dr. Rodrigo Pacheco (Fundación Ciencias para la Vida, Santiago, Chile). Knockout, wild-type and heterozygous mice littermates were obtained from heterozygous × heterozygous mattings and genotyped upon weaning with primers described in Table 1. Mice were group-housed (four-five mice/cage) in the animal facility of the Pontificia Universidad Católica de Chile and maintained in a 12/12 h inverted light/dark cycle (light on at 10.00 pm) in stable conditions of temperature (24°C), with food and water available *ad libitum*. All experimental procedures were carried out in accordance with the bioethical rules defined by the Bioethical Committee of the Pontificia Universidad Católica de Chile and the Bioethical Committee of the Consejo Nacional de Ciencia y Tecnología (CONICYT). All procedures were conducted to reduce the number of mice used when possible and to reduce their level of pain and discomfort as much as possible.

**Table 1:**
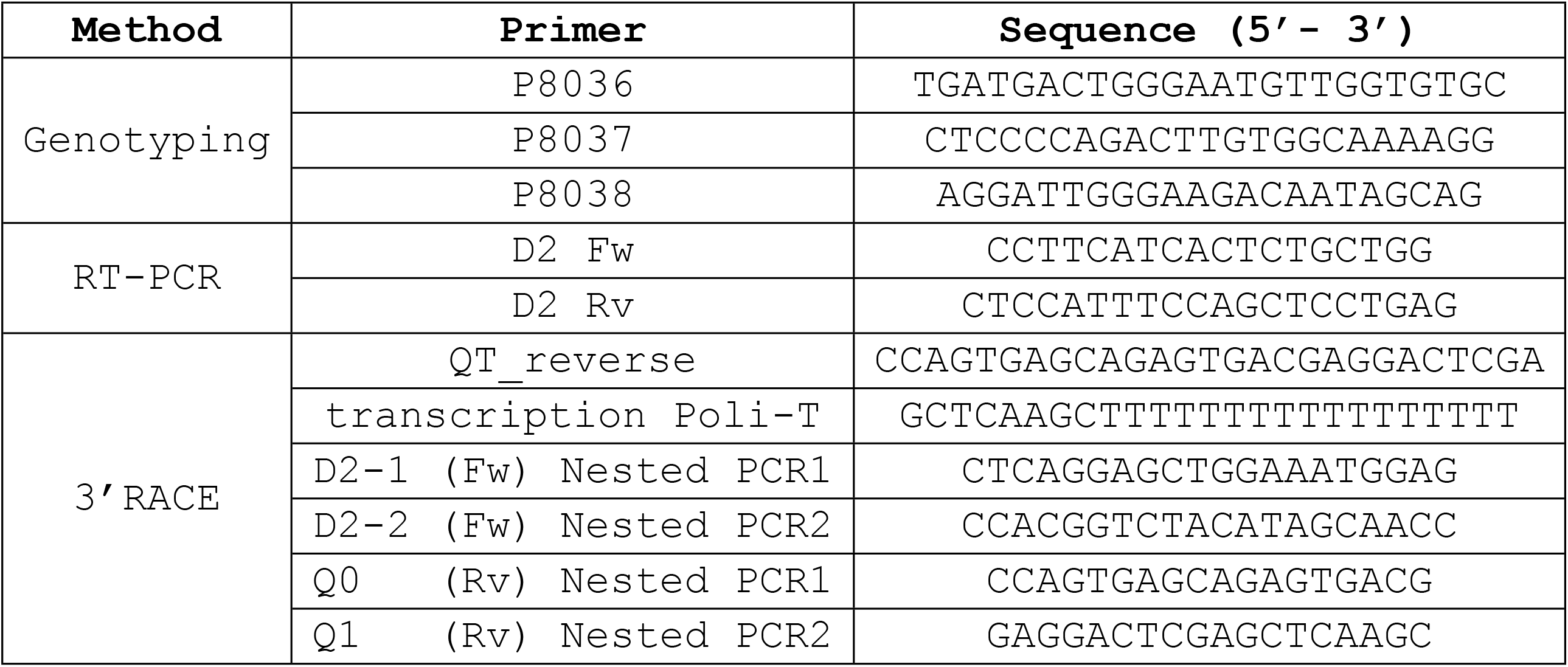
Primer sequences

### Horizontal locomotor activity

Locomotor activity was quantified as we have described [13] and adapted for mice by the use of plexiglass adapters. Briefly, mice were injected intraperitoneally (ip) two consecutive days with saline and on the third day with QNP (5mg/kg, ip). Immediately after each injection, mice were transferred individually to test cages (15×47×26 cm) equipped with two pairs of infrared lights connected to a counting device. During 90 minutes, locomotor activity was counted only when both infrared light beams were interrupted consecutively.

### Bioinformatics research

Mouse D2R gene was analyzed with the UCSC genome browser [14] using the GRCm38/mm10 assembly of *Mus musculus* genome (December 2011). We performed an *in silico* analysis of restriction cleavage sites used in the generation of the D2R functional knockout [11] and an *in silico* PCR analysis with the primers used for genotyping these mice (Table 1). The coding sequence of the mouse D2R protein and its transmembrane segments were identified using the Ensembl project [15].

### RT-PCR and 3’RACE

Total RNA extraction of mice striatum was performed with Trizol® reagent method. cDNA was generated using MMLV reverse transcriptase (Invitrogen). The RT-PCR was performed using the primers listed in Table 1. Two μg of total RNA of mouse striatum were subjected to a 3’RACE using the protocol described by Scotto-Lavino and collaborators [16]. The obtained cDNA was subjected to nested touchdown PCR using specific primers for D2R mRNA (Table 1). The resulting PCR products were purified with GeneJET™ gel extraction Kit (Thermo Scientific) and cloned into pGEM-T easy vector (Promega) for sequencing. The sequences were aligned to *Mus musculus* build (mm10) using BLAT.

### *In vivo* brain Microdialysis

Mice were deeply anesthetized with ketamine-xylazine-acepromazine (50-5-1 mg/kg, ip) and placed in a stereotaxic apparatus (Stoelting, Wood Dale, IL). Additional doses of anesthesia were administered as required to maintain suppression of limb compression withdrawal reflex. Body temperature was sustained by a thermostatically controlled heating pad. Microdialysis procedure was carried out essentially as we have described [17]. Briefly, a concentric 2 mm microdialysis probe (CMA 11, CMA Microdialysis AB, Solna, Sweden) was lowered crossing the dorsal striatum using the following coordinates respect to Bregma: AP: +1.2 mm, L: 1.3 mm and 5 mm under dura (Paxinos and Franklin, 2001). At the end of the experiments, the mice were transcardially perfused with 4% (weight/vol) paraformaldehyde in 0.1 M sodium phosphate-buffered saline (PBS, pH 7.4). Brains were rapidly removed, post-fixed overnight in the same solution at 4°C, and then dehydrated in 20% of sucrose solution for two days at 4°C. Thirty-μm thick sections were cut on a cryostat (Leica). Sections were collected in 0.1 M PBS and stored at 4°C until processed for probe placement verification. Brain regions were identified using a mouse brain atlas [18], and only sections including dorsal striatum (antero-posterior distance from bregma, about 1.18-0.98mm) were selected for analysis.

### Analysis of dialysate samples

Dopamine was quantified as we have described [13]. Briefly, 5 μL of the sample were injected into a HPLC system (BAS America, West Lafayette, IN, USA) consisting of a micropump (series 200, Perkin Elmer), a Unijet microbore C-18 reverse phase column (BAS), and an amperometric detector (LC4C, BAS). The mobile phase consisted of 0.1 M NaH_2_PO_4_, 0.8 mM EDTA, 1.2 mM 1-octanesulfonic acid and 4% CH_3_CN at pH 3.0, and it was pumped at 80 μL/min at room temperature. The potential of the amperometric detector was 650 mV.

Glutamate and GABA were quantified as we have described [19]. Briefly, 10 μL of the sample were mixed with 10 μL of bi-distilled water, 4 μL of borate buffer (pH 10.8) and 4 μL of florigenic reagent (20 mg of orthophthaldehyde and 10 μL of β-mercaptoethanol in 5 mL of ethanol). Ninety seconds later, the samples were injected into a HPLC system with the following configuration: quaternary gradient pump (Jasco Co. Ltd., Tokio, Japan), a C-18 reverse phase column (Kromasil®; Eka Chemicals, Bohus, Sweden), and a fluorescence detector (Jasco Co. Ltd.). The mobile phase was 0.1 M NaH_2_PO_4_ and CH_3_CN 14.5% (pH 5.7) pumped for 1 min; a continuous gradient of CH_3_CN (14.5-39.5%) pumped during the next 5 minutes, to return gradually to the initial condition of CH_3_CN 14.5% (pH 5.7) in the next 20 minutes.

### Immunofluorescence

For figure 3A-C, D2R immunofluorescence was performed in paraformaldehyde-fixed brain tissue as we have described [17]. Briefly, brain slices were incubated with goat anti-D2R polyclonal antibody (1:100, N-19 sc-7522, Santa Cruz Biotechnology) overnight at 4°C, followed by anti-goat Alexa 488 (1:500, Invitrogen). For Figures 3D-I, D2R immunofluorescence was performed in 50 μm free-floating slices. For antigen retrieval, the slices were treated in a solution of 1% NaOH + 1% H_2_O_2_ for 20 minutes at room temperature, then the samples were submerged in 0.3% of glycine in PBS for 10 minutes at room temperature, followed by the blocking in 3% BSA + 0.3% triton-100 for 1 hour at room temperature. The samples were incubated with a goat anti-D2R polyclonal antibody (1:100, Santa Cruz Biotechnology) or rabbit anti-D2R polyclonal antibody (1:100, AB5084P Merck Millipore) overnight at 4°C in blocking solution. After washing, the slices were incubated with anti-goat or anti-rabbit Alexa 488 (1:500, Invitrogen) in blocking solution for 1 hour at room temperature. DNA was stained by incubating the slices with TOTO 3-iodide (Invitrogen).

Immunofluorescent detection of Nur77 was performed as we have described [20]. Free-floating brain sections were incubated for 25 min at 80°C in sodium citrate-buffer (10 mM sodium citrate and 0.05% (vol/vol) Tween 20, pH 6) for antigen retrieval. Then, brain slices were incubated with anti-Nur77 monoclonal E-6 antibody (1:100, sc-166166 Santa Cruz Biotechnology) for 36 hours at 4°C, followed by donkey anti-mouse Alexa 488 (1:500, Invitrogen) antibody for 90 min at room temperature. After rinsing, DNA was stained with TOTO 3-iodide (Invitrogen).

### Image analysis

Confocal microscopy and image analysis were carried out in the Advanced Unit of Microscopy of the Pontificia Universidad Católica de Chile. Double-labeled images were obtained bilaterally using sequential laser scanning confocal microscopy (Olympus). Nur77-positive cells in the dorsal striatum were counted in 350 × 350 μm area of confocal images, using Image J software (NIH). Image analyses were performed by a blind observer to D2R genotype.

### Statistical analysis

Neurotransmitters levels obtained by microdialysis and Nur77 protein expression levels were analyzed using one-way ANOVA followed by Tukey’s test. Locomotor activity was analyzed two-way ANOVA followed by Bonferroni test, and dopamine levels after QNP injection were analyzed with T-test. All statistical analyses were performed with Graphpad Prism 5.01 software.

## Results

### D2R functional knockout mice lack the exon 8 predicting a truncated protein

The D2R functional knockout mouse was generated by inserting a neomycin cassette into the genomic site corresponding to exon 7 and most of exon 8, according to the mice genome description available at that time [11]. This D2R knockout mice line is available in the Jackson laboratory (B6.129S2-Drd2tm1Low) from where it was obtained. Genotyping was performed with the mix of primers suggested by the Jackson laboratories (Table 1), which consists of a common forward primer located in intron 7 (blue, Fig. 1A) plus two reverse primers, one also located in intron 7 (purple, Fig. 1A) and the other that hybridizes with the neomycin cassette (green, Fig. 1A). As shown in figure 1B, a band of lower migration corresponding to the mutant allele was amplified from knockout and heterozygous mice, and the band corresponding to the expected amplification for the native allele was amplified from wild-type and heterozygous mice (Fig. 1B).

**Figure 1:**
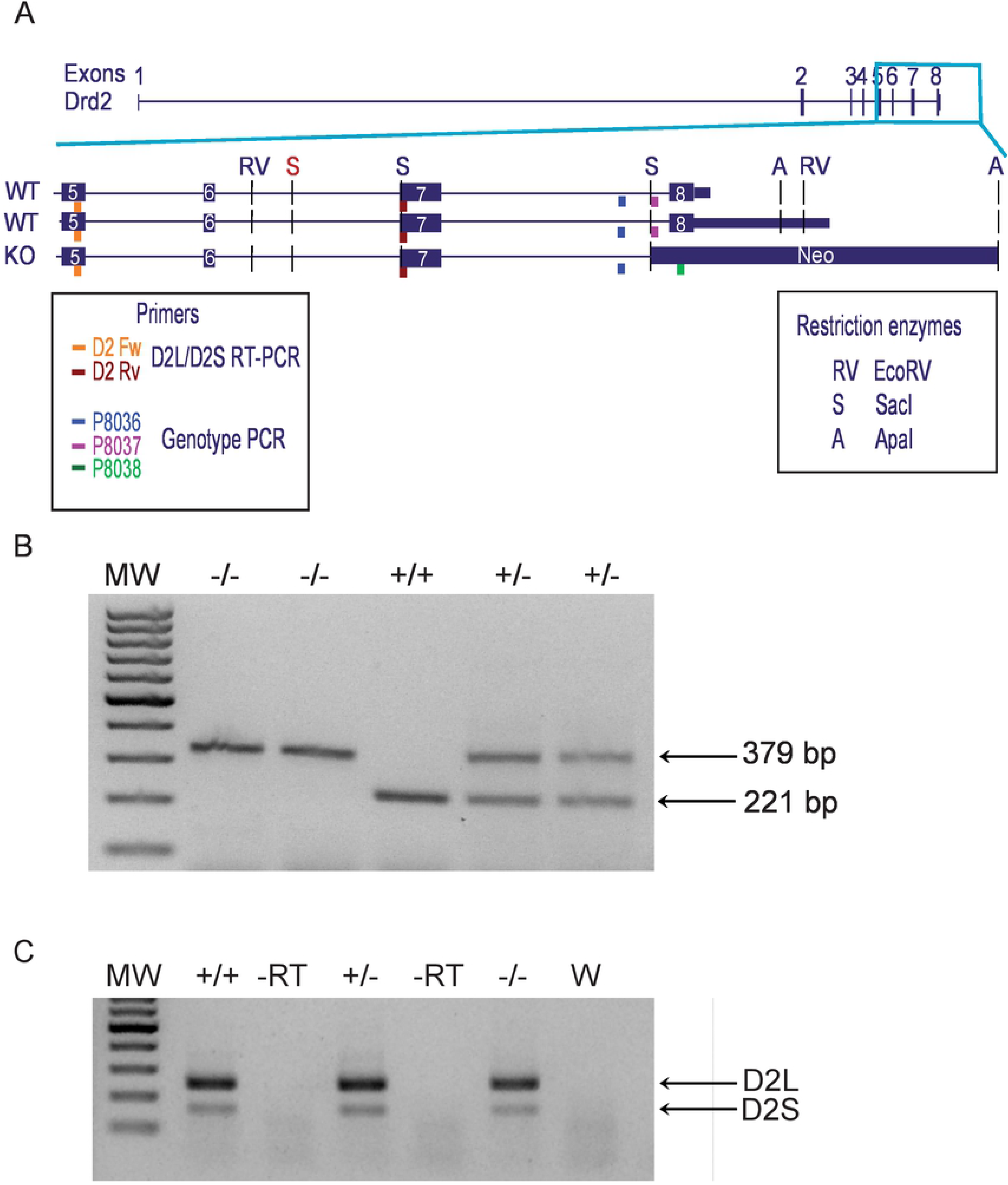
D2R functional knockout mice express mRNA transcript and lose the exon 8. (**A)** Scheme depicting mice *Drd2* gene and a zoom in fragment encompassing exons 5 to 8. Two isoforms of gene are showed with long and a short alternatives of 3’UTR. The location of Neomycin is showed in the knockout mice gene. Colored boxes show the primers used in the RT-PCR to detect D2L and D2S spliced variants, and primers used for D2R mice genotyping. Restriction enzymes cleavage sites used to generate the D2R functional knockout (Kelly et al, 1997) are indicated (in red, extra Sac1 cleavage site found). (**B)** Example of a genotyping gel of the three D2R mice genotypes. The 221 bp band correspond to wild-type (+/+) genotype and the 329 bp band correspond to knockout (−/−) genotype, heterozygous show both bands. **C)** RT-PCR products obtained from mice striatum of the three D2R genotypes amplified using primers located on exons 5 and 7 that recognize both D2L and D2S splice variants. MW: molecular weight markers, +/+ wild-type, +/− heterozygous, −/− knockout, -RT without reverse transcriptase and W PCR control with H_2_O.

After genotyping, we analyzed the expression of D2R mRNA in the striatum of mice brains. Northern analysis performed in the original paper [11] showed that heterozygous mice expressed approximately 50% of the D2R mRNA compared to wild-type, and knockout mice showed very weak expression of a smaller transcript. Therefore, we proceed to perform a diagnostic RT-PCR, with primers located in exons 5 and 7 (Table 1, Fig.1A), expecting no amplification from knockouts. Unexpectedly, RT-PCR using the reverse primer located in exon 7 amplified a product from the total mRNA obtained from the striatum of knockout mice. Also, the PCR product obtained from the striatum of knockout mice was equivalent to that of the wild-type and heterozygous in size and quantity (Fig. 1C), indicating that the D2R functional knockout mice express a D2R mRNA that includes the exon 7.

The fact that D2R functional knockout mice expressed an appreciable amount of D2R mRNA including the exon 7, prompted us to re-analysis how the *Drd2* gene was interrupted, using the most recent sequence available from the mouse genome. According to the genome finder at the University of California Santa Cruz (UCSC), the gene *Drd2* of mice has 8 exons and 7 introns, with a long and a short alternatives of 3’UTR (Fig. 1A). To identify the site where the neomycin resistance cassette was inserted, an *in silico* analysis of the position of the sequences for the restriction enzymes used in the original paper [11] was performed. As shown in figure 1A, there is an additional cutting site for the SacI enzyme, not previously noted. The SacI-ApaI cut, where the neomycin cassette was inserted, leaves exon 7 intact but eliminates exon 8 (Fig. 1A). To further prove that the D2R functional knockout mice only lacks exon 8, and not exon 7 and part of exon 8 as described in the original paper [11], we carried out a 3’RACE experiment using a forward primer located in the exon 7 and a polyT reverse primer (Table 1) to compare mature poly-A transcripts generated in the striatum of the three D2R genotypes. One main band was obtained from wild-type mice, identified as WT1 in the gel (Fig. 2A). Sequence analysis of WT1 showed that corresponds to the D2R cDNA with the longest 3’UTR (Fig. 2B). As expected, the products profile of the heterozygous mice corresponds to the sum of the products observed in the wild-type and knockout mice (Fig. 2A). Sequencing and alignment analysis of products obtained from knockout showed that the mayor product (KO1) corresponds to a mature transcript bearing the entire exon 7 but not the exon 8 of the *Drd2* gene (Fig. 2A). In addition, KO1 brings part of the last exon of an unrelated downstream gene (Ttc12) that we predicted that fuses 15 amino acids to the end of the truncated D2R (Fig. 2B and Supple Fig 1). The other minor bands (KN) correspond to truncated transcripts that have in common the loss of the exon 8 (Fig. 2A and Supple Fig 1). Altogether, these results indicate that the *Drd2* gene lacking exon 8 in the D2R functional knockout is transcribed generating stable mRNAs.

**Figure 2:**
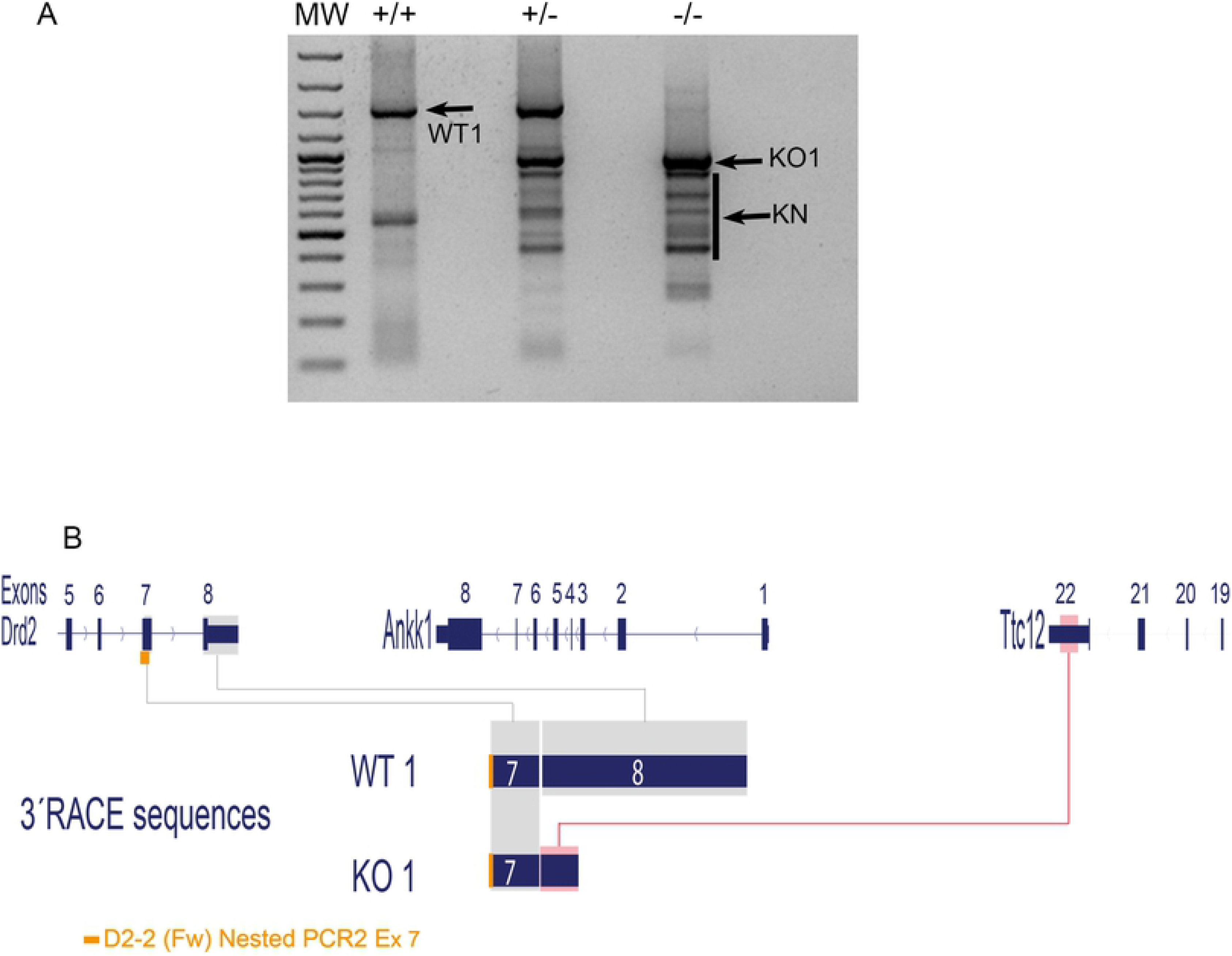
Knockout mice express different truncated transcripts that lack the exon 8. **(A)** 3’RACE gel with the generated transcripts in wild-type (+/+; WT1), heterozygous (+/−) and knockout mice (−/−; KO1, KN). The indicated bands were cloned and sequenced. **(B)** Scheme of mice *Drd2* gene as annotated in the UCSC mouse genome including downstream neighbor genes. Sequence alignment of WT1 3’RACE product, match the wild-type annotated longest D2R mRNA (+/+; WT1). Sequence alignment of KO1 3’RACE product, match with complete exon 7 of D2R plus part of the last sequence of Ttc12 gene (−/−; KO1).

### D2R functional knockout mice have a protein recognizable by anti-D2R antibodies

The fact that D2R mRNA transcripts obtained from heterozygous and knockout mice were easily detectable (Fig. 1C) suggested that the D2R-*Δe8* mRNA is stable and potentially translatable to a protein. To test this idea, we performed immunofluorescence assays in brain slices of the three D2R genotypes. Specific D2R-like immunofluorescence was observed in the striatum and nucleus accumbens of wild-type mice adult brain using a goat polyclonal (N19, Santa Cruz Biotechnology) that recognizes the N-terminal of D2R protein oriented towards the extracellular space (Fig. 3 Ai, Aii, Aiii, D) and a rabbit polyclonal (AB5084P, Millipore) that recognizes a 28-amino acids peptide localized in the 3^rd^ intracellular loop common to D2S and D2L (Fig. 3G). An inspection at higher magnifications shows D2R positive neurons in the nucleus accumbens core and shell (Fig. 3Aii, Aiii) and striatum (Fig. 3D, G), and abundant punctated mark indicative of fibers positive for D2R (Fig. 3Ai, Aii, Aiii). Significant lower D2R-like immunofluorescence was observed in knockout animals (Fig. 3Ci, Cii, Cii, E, H). Interestingly, most of the punctated mark indicative of fibers positive for D2R was lost in the D2R functional knockout while the mark for D2R in the soma of neurons in the nucleus accumbens (Fig. 3 Cii, Ciii,) and striatum (Fig. 3E, H) looks similar to the wild-type (Fig. 3 Aii, Aiii, D, G). The heterozygous mice showed similar D2R-like immunostaining to that of wild-type mice (Fig. 3 Bi, Bii, Biii). Altogether, the data indicate that a truncated D2R protein is translated from the transcripts produced in the D2R functional knockout mice.

**Figure 3:**
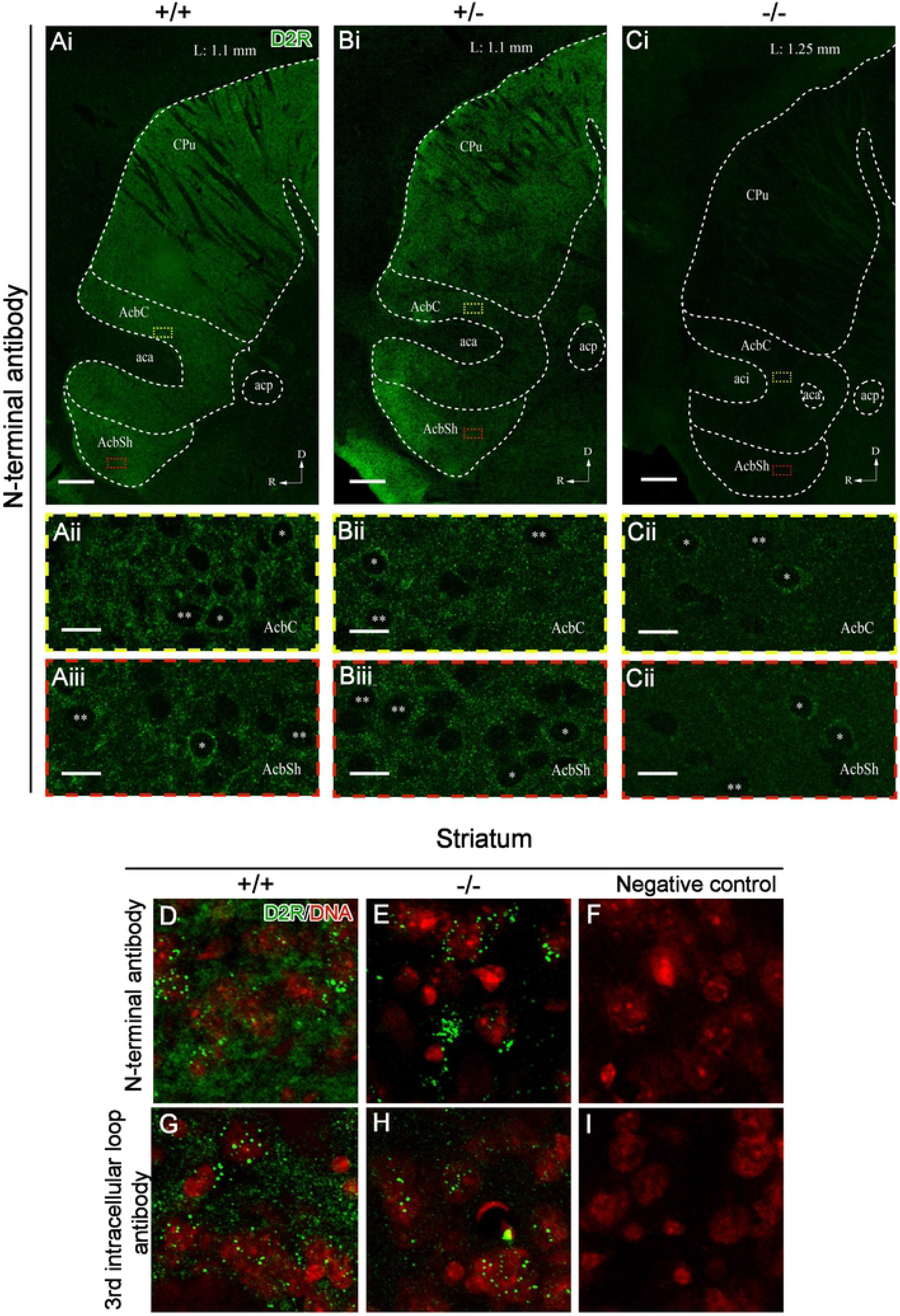
A D2R-like protein is detected in the striatum of D2R functional knockout mice. **(A-C)** Immunofluorescence for D2R in brain sagittal slices of **(A)** wild-type (+/+), **(B)** heterozygous (+/−), and **(C)** knockout (−/−) mice using a goat-anti D2R polyclonal antibody that recognizes the N-terminal region (Santa Cruz Biotechnology). (**Ai-Ci**) Sagittal view of representative reconstruction from 10X images of the striatum (CPu), nucleus accumbens core (AcbC) and shell (AcbSh); scale bar = 100μm. 100X magnification of AcbC indicated as yellow-square crop **(Aii-Cii)** and AcbSh indicated as a red-square crop **(Aiii-Ciii)**. *Positive D2R labeled cell body; **negative D2R cell body; scale bar = 10μm. aca (anterior part of anterior commissure); acp (posterior part of anterior commissure); aci (anterior commissure); R (rostral); D (dorsal); L (lateral). Immunofluorescence for D2R in the striatum of wild-type **(D, G)** and knockout **(E, H)** mice using the N-terminal recognizing antibody (**D, E**) and the 3^rd^ intracellular loop recognizing antibody (**G, H**). Negative controls without primary antibodies (**F, I**).

### Comparison of dopamine neurotransmission and locomotor activity in the three genotypes of D2R functional knockout mice

According to the 3’RACE results (Fig. 2), the putative truncated D2R protein lacks the transmembrane domains 6^th^ and 7^th^, and the C-terminal end of D2R, which are encoded in exon 8. It has been reported that D2R forms homodimers and interacts with other proteins that regulate its function and recycling [21, 22]. Thus, we wonder if the coexistence of the native D2R protein with the mutant, as it occurs in the heterozygous, could have effects different from those observed in the wild-type and knockout. To test this idea, we analyzed the locomotor activity and dopamine extracellular levels in the striatum of the three genotypes of the D2R functional knockout mice, under basal conditions and after an acute injection of the D2R agonist QNP.

Quantification of locomotor activity under basal conditions showed that the knockout mice were slower and moved less than wild-type littermates after an acute injection of saline (Fig. 4A). On the other hand, the heterozygous mice showed a basal locomotor activity slightly higher than wild-type littermates (Fig. 4A). As expected, an acute injection of QNP produced a strong significant decrease of locomotor activity in wild-type mice while the knockout mice did not respond to QNP action (Fig. 4A). The heterozygous mice showed a significant decrease of the locomotor activity similar to wild-type animals (Fig. 4A). In resume, heterozygous mice display a locomotor activity and response to an acute stimulation of D2R similar to wild-type animals.

**Figure 4:**
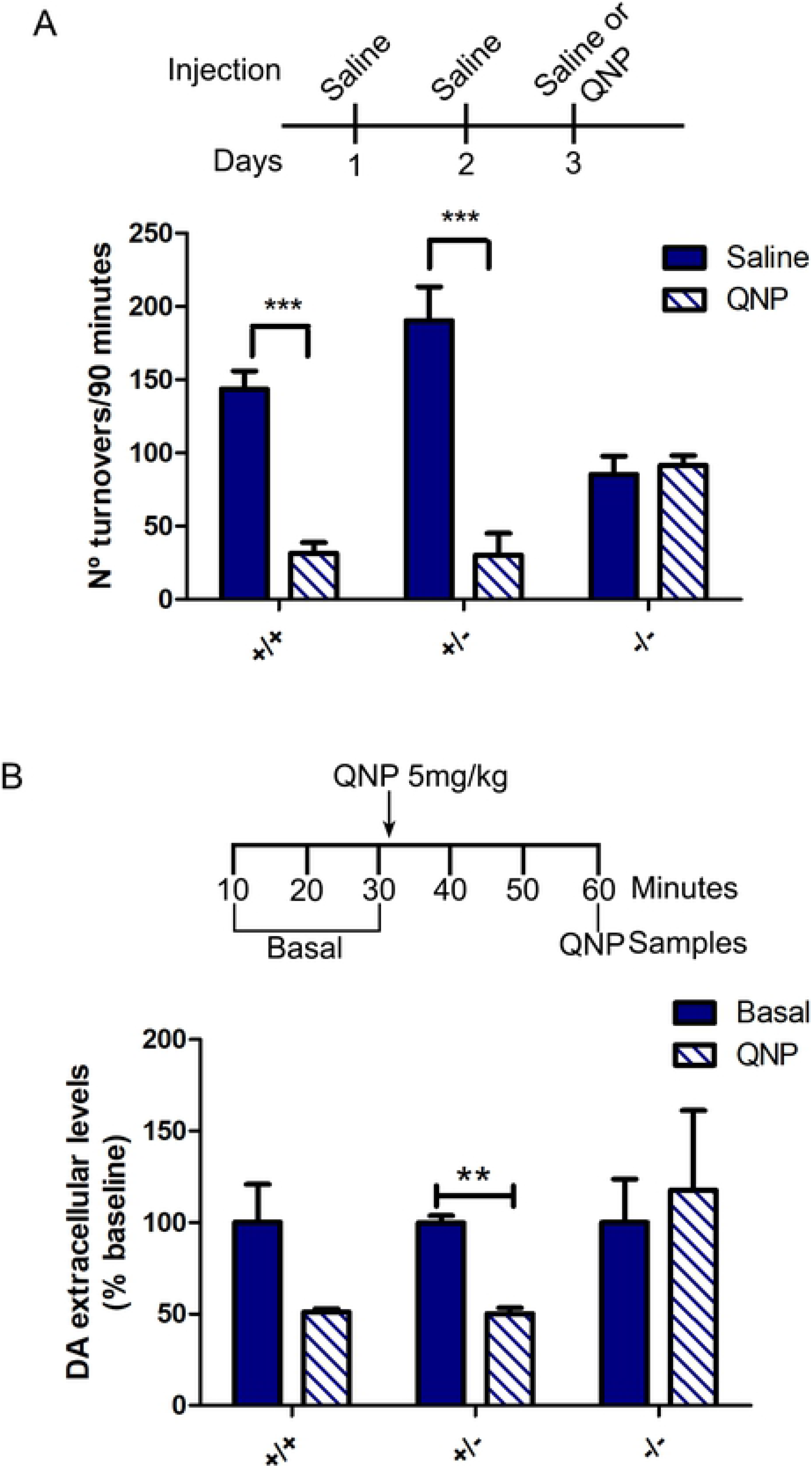
An acute injection of QNP decreases the horizontal locomotor activity in wild-type and heterozygous mice, but no in knockout D2R mice, which correlate with changes in dopamine extracellular levels in the striatum. **(A)** The three genotypes of D2R mice were injected with saline (days 1 and 2) and saline or QNP (day 3). After each injection, horizontal locomotor activity was measured during ninety minutes. Two-way ANOVA interaction: F_(2,34)_=8.631, p <0.001, treatment effect: F_(1,34)_= 28.01, p <0.01, genotype effect: F_(2,34)_= 0.7393, p >0.05. ***p< 0.001. (**B)** Microdialysis samples were collected every 10 minutes. Three samples were obtained to calculate basal extracellular levels of dopamine. Then, mice received a QNP (5 mg/kg, ip) injection, and after 30 minutes, a sample was obtained to quantify dopamine. **p<0.01.

Surprisingly, microdialysis data showed that heterozygous mice, like knockouts, have half the basal extracellular levels of dopamine in the striatum, compared to wild-type littermates (Table 2). Accordingly, the expression of Nur77, a transcription factor and immediate-early gene whose expression is a readout of dopamine neurotransmission in the striatum [23] was similarly lower in the striatum of both heterozygous and knockout mice (Supple Fig. 2), as previously reported [24].

**Table 2:** Extracellular levels of dopamine, glutamate, and GABA in the striatum of the three genotypes of functional D2R knockout mice

| Genotype/Neurotransmitter | Dopamine (fmol/μl) | Glutamate (pmol/μl) | GABA (pmol/μl) |
| --- | --- | --- | --- |
| +/+ | 0.95 ± 0.20 | 1.48 ± 0.55 | 0.024 ± 0.003 |
| +/- | 0.33 ± 0.03<sup>a</sup> | 1.45 ± 0.59 | 0.027 ± 0.002 |
| -/- | 0.36 ± 0.05<sup>b</sup> | 1.33 ± 0.25 | 0.033 ± 0.006 |

One-way ANOVA: $F(2,15) = 7.68$ , $p = 0.0051$ . Post hoc Tukey's test, +/+ vs +/- <sup>a</sup>
$p < 0.05$ and +/+ vs -/- <sup>b</sup> $p < 0.01$ .

Although, hippocampal and prefrontal glutamate inputs to the striatum are also under presynaptic D2R negative control [8], unchanged glutamate levels were observed in heterozygous and knockout mice (Table 2). Similarly, GABA extracellular levels were not significantly modified in the heterozygous and knockout mice. As expected, microdialysis data showed that the acute injection of QNP decreased the extracellular levels of dopamine in the wild-type mice while the knockout did not show changes (Fig. 4B). Notably, QNP injection induced a further 50% decrease of extracellular dopamine levels in the striatum of heterozygous mice (Fig. 4B). Therefore, heterozygous mice have similar synaptic dopamine concentration as knockouts but respond to QNP similarly to wild-types.

## Discussion

Our work shows that the D2R functional knockout is better described as a D2 functional knocking mouse, in which the *Drd2* gene has a deletion of the exon 8. This modified *Drd2ΔE8* gene is transcribed, leading to a truncated protein that lacks the last 64 amino acids, which encode transmembrane domains sixth and seventh, and the C-terminal end of D2R (Fig. 5). Our data indicate that a D2R-like protein is present in the brain of the D2R functional knockout mice.

**Figure 5:**
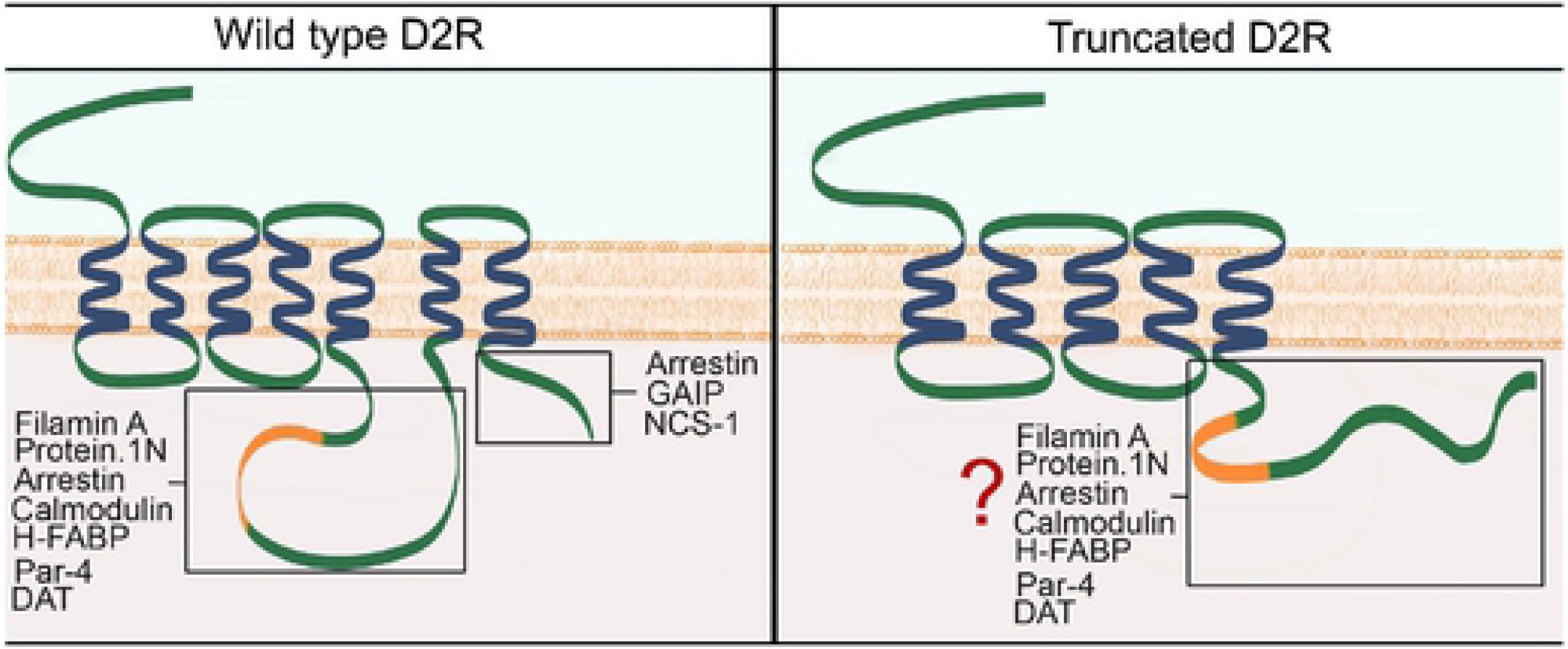
Working model of the putative truncated protein in the D2R functional knockout mice. Scheme of wild-type D2R protein and D2R truncated protein as predicted by Ensembl project. D2R interacting proteins related to trafficking, location, and cell signaling are depicted. The working model indicates that protein interaction though the C-terminal domain is lost in the D2R functional knockout while it could maintain the interaction though the 3^rd^ intracellular loop.

The comparative analysis of the locomotor activity and extracellular levels of dopamine in the three genotypes showed that the heterozygous mice share some features with wild-types while others are similar to knockout mice. Indeed, basal locomotor activity and response to QNP of heterozygous mice are indistinguishable from that of wild-types, while knockouts display lower locomotor activity and do not respond to QNP. On the other hand, the heterozygous and knockout mice have a significant similar lower extracellular dopamine levels in the striatum compared to wild-types. However, heterozygous respond to QNP, decreasing further the extracellular levels of dopamine, like the wild-types. These data suggest that lower locomotor activity in D2R functional knockout mice is not due to reduced levels of synaptic dopamine but to lack of postsynaptic D2R signaling, and the locomotor activity is the result of changes in striatal levels of dopamine rather than its absolute values.

Lower extracellular dopamine levels in the nucleus accumbens of the D2R functional knockout were previously reported [25, 26]. Work performed by Schmitz et al. (2002) indicated that the null D2R knockout had increased dopamine uptake [27]. Therefore, lower dopamine extracellular levels in D2R functional knockout and heterozygous mice could be due to higher uptake. The mixed features in behavior and dopamine neurotransmission showed by the heterozygous mice could be due to the coexistence of the truncated with the native D2R. The truncated D2R protein localized in dopaminergic axon terminals or postsynaptically in the MSN or interneurons in the striatum could opposes to the role of the native D2R protein, acting as a dominant-negative by interfering with interaction and trafficking of proteins that control extracellular levels of dopamine as the dopamine transporter [28] or postsynaptically with signaling proteins as β-arrestin [29].

Immunofluorescence data showing D2R-like immunolabeling in the striatum of the knockout mice support the presence of a D2R-like protein that was recognized by two antibodies, one aimed to the N-terminal and the other to the third intracellular loop of D2R. This truncated protein losing the sixth and seventh transmembrane segments and the C-terminal end (see a model in Fig. 5) could modify the interaction with proteins as β-arrestin, GAIP and NCS-1, which regulate the internalization of D2R [30]. Moreover, incorrect folding of the last part of the predicted truncated protein could modify the interaction between this fragment of D2R with proteins as filaminA and protein 4.1N, both related to D2R cell surface localization [31]. Although the truncated D2R protein should still interact with proteins as G_i/o_ [32] and the dopamine transporter [28], changes in localization and availability of the receptor could explain the mixed characteristic of the heterozygous mice. These characteristics make this D2R functional knockout and heterozygous mice a valuable tool to understand the interactions between D2R and proteins that regulate its function and localization.

## Acknowledgments

The authors thank Dr. Rodrigo Pacheco for kindly provide the D2R functional knockout mice. The authors appreciate the comments and suggestions made by Dr. Héctor Yarur. The authors acknowledge the services provided by UC CINBIOT Animal Facility funded by PIA CONICYT* ECM-07 *Program for Associative Research, of the Chilean National Council for Science and Technology. The authors acknowledge the services provided by the Advanced Unit of Microscopy of the Pontificia Universidad Católica de Chile.

## Supporting information

**S1 Fig. Scheme depicting mice** *Drd2* **gene and zoom in fragment encompassing exons 7 to 8.** Alignment of minor bands from the 3`RACE experiment (KN). Upper part, sequences that include exon 7 (grey box) and a part of the intron 7 (yellow box). In the lower part, sequences that include the exon 7 and also a sequence located in intron 2 (orange and green boxes) or intron 6 (red box) of Ttc12 gene.

**S2 Fig. Lower Nur77 expression in the striatum of D2R functional heterozygous and knockout mice**. (A-L) Confocal sections of the striatum showing immunofluorescence for Nur77 (green) in wild-type (A-D), heterozygous (E-H), and knockout (I-L) mice. Cell nuclei were stained with TOTO3 (red). (D, H, L) zoomed view of the area indicated by a square in the merge panel. Scale bar: 100 μm. (M) Quantification of the number of Nur77 positive cells in the striatum of each genotype of D2R mice. Data are means ± SEM. One-way ANOVA: F(2,8) = 12.16, p = 0.0038. Post hoc Tukey’s test, +/+ vs +/− **p<0.01 and +/+ vs −/− *p<0.05.

## References

1. Giros B, Sokoloff P, Martres MP, Riou JF, Emorine LJ, Schwartz JC. Alternative splicing directs the expression of two D2 dopamine receptor isoforms. Nature. 1989;342(6252):923–6. Epub 1989/12/21. doi: 10.1038/342923a0. PubMed PMID: 2531847.

2. Monsma FJ, Jr., McVittie LD, Gerfen CR, Mahan LC, Sibley DR. Multiple D2 dopamine receptors produced by alternative RNA splicing. Nature. 1989;342(6252):926–9. Epub 1989/12/21. doi: 10.1038/342926a0. PubMed PMID: 2480527.

3. De Mei C, Ramos M, Iitaka C, Borrelli E. Getting specialized: presynaptic and postsynaptic dopamine D2 receptors. Curr Opin Pharmacol. 2009;9(1):53–8. Epub 2009/01/14. doi: 10.1016/j.coph.2008.12.002.. PubMed PMID: 19138563 PubMed Central PMCID: PMCPMC2710814.

4. Radl D, Chiacchiaretta M, Lewis RG, Brami-Cherrier K, Arcuri L, Borrelli E. Differential regulation of striatal motor behavior and related cellular responses by dopamine D2L and D2S isoforms. Proc Natl Acad Sci U S A. 2018;115(1):198–203. Epub 2017/12/20. doi: 10.1073/pnas.1717194115.. PubMed PMID: 29255027 PubMed Central PMCID: PMCPMC5776825.

5. Deng YP, Lei WL, Reiner A. Differential perikaryal localization in rats of D1 and D2 dopamine receptors on striatal projection neuron types identified by retrograde labeling. J Chem Neuroanat. 2006;32(2-4):101–16. Epub 2006/08/18. doi: 10.1016/j.jchemneu.2006.07.001. PubMed PMID: 16914290.

6. Bertran-Gonzalez J, Bosch C, Maroteaux M, Matamales M, Herve D, Valjent E, et al. Opposing patterns of signaling activation in dopamine D1 and D2 receptor-expressing striatal neurons in response to cocaine and haloperidol. J Neurosci. 2008;28(22):5671–85. Epub 2008/05/30. doi: 10.1523/JNEUROSCI.1039-08.2008. PubMed PMID: 18509028; PubMed Central PMCID: PMCPMC6670792.

7. Smith Y, Bevan MD, Shink E, Bolam JP. Microcircuitry of the direct and indirect pathways of the basal ganglia. Neuroscience. 1998;86(2):353–87. Epub 1999/01/09. doi: 10.1016/s0306-4522(98)00004-9. PubMed PMID: 9881853.

8. Sesack SR, Grace AA. Cortico-Basal Ganglia reward network: microcircuitry. Neuropsychopharmacology. 2010;35(1):27–47. Epub 2009/08/14. doi: 10.1038/npp.2009.93. PubMed PMID: 19675534; PubMed Central PMCID: PMCPMC2879005.

9. Missale C, Nash SR, Robinson SW, Jaber M, Caron MG. Dopamine receptors: from structure to function. Physiol Rev. 1998;78(1):189–225. Epub 1998/02/11. doi: 10.1152/physrev.1998.78.1.189. PubMed PMID: 9457173.

10. Baik JH, Picetti R, Saiardi A, Thiriet G, Dierich A, Depaulis A, et al. Parkinsonian-like locomotor impairment in mice lacking dopamine D2 receptors. Nature. 1995;377(6548):424–8. Epub 1995/10/05. doi: 10.1038/377424a0. PubMed PMID: 7566118.

11. Kelly MA, Rubinstein M, Asa SL, Zhang G, Saez C, Bunzow JR, et al. Pituitary lactotroph hyperplasia and chronic hyperprolactinemia in dopamine D2 receptor-deficient mice. Neuron. 1997;19(1):103–13. Epub 1997/07/01. doi: 10.1016/s0896-6273(00)80351-7. PubMed PMID: 9247267.

12. Jung MY, Skryabin BV, Arai M, Abbondanzo S, Fu D, Brosius J, et al. Potentiation of the D2 mutant motor phenotype in mice lacking dopamine D2 and D3 receptors. Neuroscience. 1999;91(3):911–24. Epub 1999/07/03. doi: 10.1016/s0306-4522(98)00705-2. PubMed PMID: 10391470.

13. Escobar AP, Cornejo FA, Olivares-Costa M, Gonzalez M, Fuentealba JA, Gysling K, et al. Reduced dopamine and glutamate neurotransmission in the nucleus accumbens of quinpirole-sensitized rats hints at inhibitory D2 autoreceptor function. J Neurochem. 2015;134(6):1081–90. Epub 2015/06/27. doi: 10.1111/jnc.13209. PubMed PMID: 26112331.

14. Kent WJ, Sugnet CW, Furey TS, Roskin KM, Pringle TH, Zahler AM, et al. The human genome browser at UCSC. Genome Res. 2002;12(6):996–1006. Epub 2002/06/05. doi: 10.1101/gr.229102.. PubMed PMID: 12045153 PubMed Central PMCID: PMCPMC186604.

15. Yates A, Akanni W, Amode MR, Barrell D, Billis K, Carvalho-Silva D, et al. Ensembl 2016. Nucleic Acids Res. 2016;44(D1):D710–6. Epub 2015/12/22. doi: 10.1093/nar/gkv1157.. PubMed PMID: 26687719 PubMed Central PMCID: PMCPMC4702834.

16. Scotto-Lavino E, Du G, Frohman MA. 3’ end cDNA amplification using classic RACE. Nat Protoc. 2006;1(6):2742–5. Epub 2007/04/05. doi: 10.1038/nprot.2006.481. PubMed PMID: 17406530.

17. Escobar AP, Gonzalez MP, Meza RC, Noches V, Henny P, Gysling K, et al. Mechanisms of Kappa Opioid Receptor Potentiation of Dopamine D2 Receptor Function in Quinpirole-Induced Locomotor Sensitization in Rats. Int J Neuropsychopharmacol. 2017;20(8):660–9. Epub 2017/05/23. doi: 10.1093/ijnp/pyx042.. PubMed PMID: 28531297 PubMed Central PMCID: PMCPMC5569963.

18. Paxinos G, Franklin KBJ. The mouse brain in stereotaxic coordinates. Academic Press. 2001

19. Sotomayor-Zarate R, Araya KA, Pereira P, Blanco E, Quiroz G, Pozo S, et al. Activation of GABA-B receptors induced by systemic amphetamine abolishes dopamine release in the rat lateral septum. J Neurochem. 2010;114(6):1678–86. Epub 2010/06/30. doi: 10.1111/j.1471-4159.2010.06877.x. PubMed PMID: 20584106.

20. Sanchez N, Coura R, Engmann O, Marion-Poll L, Longueville S, Herve D, et al. Haloperidol-induced Nur77 expression in striatopallidal neurons is under the control of protein phosphatase 1 regulation by DARPP-32. Neuropharmacology. 2014;79:559–66. Epub 2014/01/21. doi: 10.1016/j.neuropharm.2014.01.008. PubMed PMID: 24440754.

21. Lee SP, Xie Z, Varghese G, Nguyen T, O’Dowd BF, George SR. Oligomerization of dopamine and serotonin receptors. Neuropsychopharmacology. 2000;23(4 Suppl):S32–40. Epub 2000/09/29. doi: 10.1016/S0893-133X(00)00155-X. PubMed PMID: 11008065.

22. Fuxe K, Borroto-Escuela DO, Marcellino D, Romero-Fernandez W, Frankowska M, Guidolin D, et al. GPCR heteromers and their allosteric receptor-receptor interactions. Curr Med Chem. 2012;19(3):356–63. Epub 2012/02/18. PubMed PMID: 22335512.

23. Campos-Melo D, Galleguillos D, Sanchez N, Gysling K, Andres ME. Nur transcription factors in stress and addiction. Front Mol Neurosci. 2013;6:44. Epub 2013/12/19. doi: 10.3389/fnmol.2013.00044.. PubMed PMID: 24348325 PubMed Central PMCID: PMCPMC3844937

24. An JJ, Bae MH, Cho SR, Lee SH, Choi SH, Lee BH, et al. Altered GABAergic neurotransmission in mice lacking dopamine D2 receptors. Mol Cell Neurosci. 2004;25(4):732–41. Epub 2004/04/15. doi: 10.1016/j.mcn.2003.12.010. PubMed PMID: 15080900.

25. Zapata A, Shippenberg TS. Lack of functional D2 receptors prevents the effects of the D3-preferring agonist (+)-PD 128907 on dialysate dopamine levels. Neuropharmacology. 2005;48(1):43–50. Epub 2004/12/25. doi: 10.1016/j.neuropharm.2004.09.003. PubMed PMID: 15617726.

26. Fan X, Xu M, Hess EJ. D2 dopamine receptor subtype-mediated hyperactivity and amphetamine responses in a model of ADHD. Neurobiol Dis. 2010;37(1):228–36. Epub 2009/10/21. doi: 10.1016/j.nbd.2009.10.009.. PubMed PMID: 19840852 PubMed Central PMCID: PMCPMC2839459.

27. Schmitz Y, Schmauss C, Sulzer D. Altered dopamine release and uptake kinetics in mice lacking D2 receptors. J Neurosci. 2002;22(18):8002–9. Epub 2002/09/12. PubMed PMID: 12223553.

28. Lee FJ, Pei L, Moszczynska A, Vukusic B, Fletcher PJ, Liu F. Dopamine transporter cell surface localization facilitated by a direct interaction with the dopamine D2 receptor. EMBO J. 2007;26(8):2127–36. Epub 2007/03/24. doi: 10.1038/sj.emboj.7601656.. PubMed PMID: 17380124 PubMed Central PMCID: PMCPMC1852782.

29. Beaulieu JM, Sotnikova TD, Marion S, Lefkowitz RJ, Gainetdinov RR, Caron MG. An Akt/beta-arrestin 2/PP2A signaling complex mediates dopaminergic neurotransmission and behavior. Cell. 2005;122(2):261–73. Epub 2005/07/30. doi: 10.1016/j.cell.2005.05.012. PubMed PMID: 16051150.

30. Fukunaga K, Shioda N. Novel dopamine D2 receptor signaling through proteins interacting with the third cytoplasmic loop. Mol Neurobiol. 2012;45(1):144–52. Epub 2011/12/21. doi: 10.1007/s12035-011-8227-8. PubMed PMID: 22183739.

31. Kabbani N, Levenson R. A proteomic approach to receptor signaling: molecular mechanisms and therapeutic implications derived from discovery of the dopamine D2 receptor signalplex. Eur J Pharmacol. 2007;572(2-3):83–93. Epub 2007/07/31. doi: 10.1016/j.ejphar.2007.06.059. PubMed PMID: 17662712.

32. Sidhu A, Niznik HB. Coupling of dopamine receptor subtypes to multiple and diverse G proteins. Int J Dev Neurosci. 2000;18(7):669–77. Epub 2000/09/09. PubMed PMID: 10978845.

